# Genetic characterization of the complete genome of a strain of Chikungunya virus circulating in Brazil: a strategy for the surveillance and control of Arboviral diseases with epidemic potential in Latin America

**DOI:** 10.1101/2024.07.21.602720

**Authors:** Rafael Guillermo Villarreal Julio, David Pabón, Marcela Palacio, Juan Zapata, Juan Sepúlveda, Santiago Zea, Andrés Patino

## Abstract

*Chikungunya* virus (CHIKV) is a positive-sense single-stranded RNA virus that belongs to the Alphavirus genus of the Togaviridae family. It is mainly transmitted by mosquitoes *Aedes aegypti and albopictus*. Its genome encodes four non-structural proteins (NSP 1-4 encoded at the 5’ end) and three structural proteins (C, E1 and E2 structural proteins encoded at the 3’ end). Four lineages of this virus have been identified, namely the lineages from 1) West Africa, 2) East, Central and South Africa (ECSA), 3) Asia (AL) and 4) Indian Ocean (IOL). CHIKV is an endemic arbovirus circulating in 51 countries in the Americas. The clinical manifestations attributed to it are: high fever, rash, myalgia and episodes of arthralgia, which consequently lead to chronic pain and disability, especially in the joints. Whole genome sequencing of the *Chikungunya* virus is essential to understand its biology, evolution and spread, and to develop effective prevention, diagnosis and treatment strategies. This information is essential to combat the disease and minimize its impact on public health. For these reasons, the complete genome of the *Chikungunya* virus BR33, identified in the northeastern city of Recife, in the state of Pernambuco, Brazil, was sequenced. The genome has a size of 11,601 nucleotides and fragments that code for two polyproteins.

A phylogenic analysis was performed indicating that the recent Brazilian strain of CHIKV belongs to the East, Central and Southern Africa (ECSA) lineage. This phylogenetic identification is important because this particular genotype has been associated with greater damage and clinical severity, in addition to the fact that this ECSA lineage has a wide range of genomic diversity.

Until 2016, CHIKV was directly associated with travel and transmission was limited. Subsequently, in the State of Pernambuco, Brazil, the introduction of a new ECSA lineage was identified, which corresponds to the one identified in this study and which generated the largest outbreak. New CHIKV outbreaks are very likely in the near future due to the abundance of competent vectors in Brazil and a susceptible population, exposing more than 11 million inhabitants to an increasing risk of infection.

## INTRODUCTION

*Chikungunya* virus (CHIKV) is an arbovirus that belongs to the Togaviridae family, within the genus *Alphavirus*. The CHIKV virus was first isolated during an outbreak in Tanzania in 1952 (de Souza WM et al. 2024).

The transmission of the CHIKV occurs mainly through the bite of mosquitoes of the genus *Aedes*, especially *Aedes aegypti*, although it has also been observed to a lesser extent in *Aedes albopictus* (de Souza WM et al. 2024). In addition to transmission by mosquitoes, vertical transmission from mother to child has been documented as another route of infection. (Carrillo et al. 2023; Nunes et al. 2015). In humans, *Chikungunya* manifests itself with symptoms such as arthralgia, high and acute fever, myalgia, headache, conjunctivitis and, in many cases, rash. However, this disease can also result in prolonged polyarthralgia and neurological complications, which can even lead to fatal outcomes. Other authors include the following signs and symptoms, Alguridi et al. 2023 (myocarditis, in vulnerable patients such as newborns and the elderly it can cause eye diseases, pneumonia and hemorrhages. In maternal cases it can cause natal and neonatal). Jiang Ping et al. 2023 includes information on the report of meningoencephalitis as a possible complication. CHIKV infection is a major public health problem in tropical and subtropical regions. (Hakim & Aman, 2022; Nunes et al. 2015). The impact of *Chikungunya* is exacerbated by climate change and human migration, which provide a favorable environment for the expansion of Aedes mosquitoes (de Souza WM et al. 2024).

Despite posing a significant threat to public health, the reporting of *Chikungunya* cases is hampered by the circulation of other mosquito-borne viruses, such as dengue (DENV) and Zika (ZIKV), which present similar clinical symptoms. This makes syndromic surveillance the main source of detection (de Souza WM et al. 2024). The disabling nature of *Chikungunya* has generated a considerable economic burden and has led to the collapse of health systems in various regions. Local transmission of CHIKV has been documented in more than 100 countries and territories in Asia, Africa, Europe and the Americas (de Souza WM et al. 2024).

According to the World Health Organization (WHO), between December 2013 and 2017, the Caribbean islands and the Americas recorded >2.5 million suspected and confirmed cases (de Souza WM et al. 2024). Between December 2013 and June 2023, 3,684,554 cases of *Chikungunya* (suspected and laboratory confirmed) were reported in 50 countries or territories in the Americas. Between 2014 and 2015, most territories or countries with *Chikungunya* outbreaks in the Latin Caribbean reported one or two annual epidemic waves followed by a period with lower incidence or no cases (Pan American Health Organization, 2014; Cunha, M. S et al.2020).

Between December 2013 and June 2023, a total of 3,684,554 cases of *Chikungunya* (suspected and laboratory confirmed) were reported in 50 countries or territories of the American continent (de Souza WM et al. 2024). This disease has had a significant impact on public health in the region. During 2014 and 2015, most countries and territories in the Latin Caribbean experienced *Chikungunya* outbreaks, characterized by one or two annual epidemic waves followed by periods with lower incidence or absence of cases. In 2014, the most severe epidemics were concentrated in the Latin Caribbean, followed by the non-Latin Caribbean. Subsequently, in 2015, the Central American and Andean regions became the main foci of the disease (de Souza WM et al. 2024). Brazil, the largest and most populous country in Latin America, presents favorable conditions for the proliferation of the *Chikungunya* virus: a large susceptible population, a favorable climate and abundant mosquitoes *Aedes aegypti*. Since its introduction in 2014, the virus has circulated locally in the country, with the first cases concentrated mainly in the northeast region. As of 2016, Brazil has become the epicenter of *Chikungunya* epidemics in the Americas, recording 1,659,167 cases, the highest number reported in the region. Unlike other countries, Brazil has experienced annual *Chikungunya* epidemics since its introduction (de Souza WM et al. 2024).

In recent decades, Colombia has faced epidemics of various emerging arboviruses, including dengue, Venezuelan equine encephalitis, *Chikungunya* and Zika, which has generated important public health problems (de Souza WM et al. 2024).

According to the Pan American Health Organization, Colombia was ranked as the third country with the highest number of *Chikungunya* cases between 2014 and 2015, after Brazil and the Dominican Republic (de Souza WM et al. 2024).

Colombia is a hyperendemic country for arboviruses due to the widespread presence of the Aedes aegypti and Aedes albopictus mosquitoes (Caceres et al. 2024). These vectors are responsible for transmitting various arboviruses, including dengue, *Chikungunya*, and Zika (Madewell., 2020). One of the important challenges in the management of these diseases is the lack of virus-specific diagnostic tests and late detection of infections. This diagnostic ambiguity often leads healthcare workers to misdiagnose *Chikungunya* and Zika as dengue. Consequently, public healthcare funds allocated for dengue management are redirected to *Chikungunya* and Zika, compromising the accuracy of epidemiological data on dengue and making it difficult for public health systems to manage epidemic outbreaks (Cáceres et al. al. 2024).

The rapid spread of *Chikungunya* is associated with various factors, including the high mobility of people and goods.

According to the National Institute of Health (INS) of Colombia, the first cases reported in the country were imported from neighboring countries such as Venezuela, the Dominican Republic and Panama. Its distinctive feature is its envelope, which contains a positive-sense single-stranded RNA genome with a length of 11.8 kb.

Phylogenetic analyzes have revealed the existence of three distinct CHIKV genotypes: West African (WA), African Eastern/Central/South Africa (ECSA) and Asia/Caribbean.

2005-2006, a new descendant of the ECSA lineage, known as the Indian Ocean lineage (IOL), was described, which developed a greater affinity for the Aedes albopictus vector due to mutations such as E1-A226V, E2-I211T, E1-T98A and E2-L210Q (de Souza WM et al. 2024).

He genotype Asia/Caribbean is endemic/epidemic mostly between human populations. ECSA initially occurred isolated in Africa, however, some lineages of it have expanded their range, causing outbreaks in tropical human populations. West Africa occurs predominantly in the west of the African continent, where it is transmitted mainly between non-human primates, with sporadic infections to humans.

Numerous CHIKV outbreaks have occurred globally due to the introduction of genotypes previously confined to certain regions into new areas with large human populations that had not been exposed before, or into areas where other genotypes were already circulating.

This spread has allowed CHIKV to spread globally, radically changing the locations of epidemics.

To the best of our knowledge, these three different phylogenetic groups of CHIKV have different antigenic properties: the East, Central and South African (ECSA) genotypes. The first cases of indigenous CHIKV in Brazil were confirmed in the city of Oiapoque, in the state of Amapá, back in September 2014, where two genotypes of CHIKV, ECSA and Asian, were identified (Alguridi et al. 2023; Nunes et al. 2015; Volk et al., 2010).

The present study sequenced the complete genome of the *Chikungunya* virus, genotype BR33 isolated from a pregnant woman identified in the northeastern city of Recife, in the state of Pernambuco, Brazil in March 2016, the sample was taken from a pregnant woman where it was initially suspected that was infected with Zika. The diagnosis of CHIKV was made by RT-PCR, designed and standardized by the Gehrke Laboratory at the Massachusetts Institute of Technology (MIT), in which a section of the E1 gene was amplified.

## METHODS

The CHIKV strain was propagated in Vero-E6 cells (Vero) that were maintained in minimal essential medium (MEM) (Sigma), supplemented with 5% to 10% fetal bovine serum and antibiotics at 37° in 5%CO2. C6/36 cells were maintained in Leibovitz’s L-15 medium (Invitrogen, Carlsbad, California, USA) supplemented with 10% fetal bovine serum, antibiotics, and 1% TPB (Sigma, St. Louis, Missouri, USA) at 32°C. collected 7 days after infection or after showing a cytopathogenic effect (Ang et al. 2016; Miller et al. 2018).

All experiments were carried out in a biosafety level 3 laboratory. RNA extraction was performed using the QIAamp Viral RNA mini kit from Qiagen, following the instructions in the instructions. Whole-genome Novo sequencing was performed on an Illumina HiSeq 2500 system (Conteville et al. 2016; Xf et al. 2023) using the Trinity RNA seq assembly method (Haas et al. 2013; Hartline et al. 2023), and version r2013-02-25. Sequencing and assembly were carried out at the Massachusetts Institute of Technology (MIT) facilities.

The characterization was performed using the ViPR database (www.viprbrc.org). The function was generated by the InterPro program (http://www.ebi.ac.uk/interpro/protein/A0A192GR82). He analysis Phylogenetic analysis was performed using Molecular Evolutionary Genetics Analysis software. (MEGA) version 7.0 (7-17) with the neighbor-joining tree method.

## RESULTS

The complete genome sequence was named *Chikungunya* virus isolate BR33 and was submitted to NCBI GenBank, with the accession number: KX228391.1. https://www.ncbi.nlm.nih.gov/nuccore/1036639523

The genome has a size of 11,601 nucleotides, and two open reading frames are evident that code for two precursor polyproteins, these are the structures. Natural and non-structural polyprotein of *Chikungunya*. The first is a nonstructural polyprotein ranging from nucleotide position 80 to 7504. The product is called CHIKVgp1 and its ID is ANK58564.1 (Fig. 1). The second structural polyprotein runs from nucleotide position 7570 to 11,316, the product is called CHIKVgp2 and its ID is ANK58565.1 (Table 1). The functional and structural features are described in detail (Fig. 1-4). In addition, phylogenetic classification of the *Chikungunya* virus isolate BR33 was performed (Fig. 5).

**Table 1.**
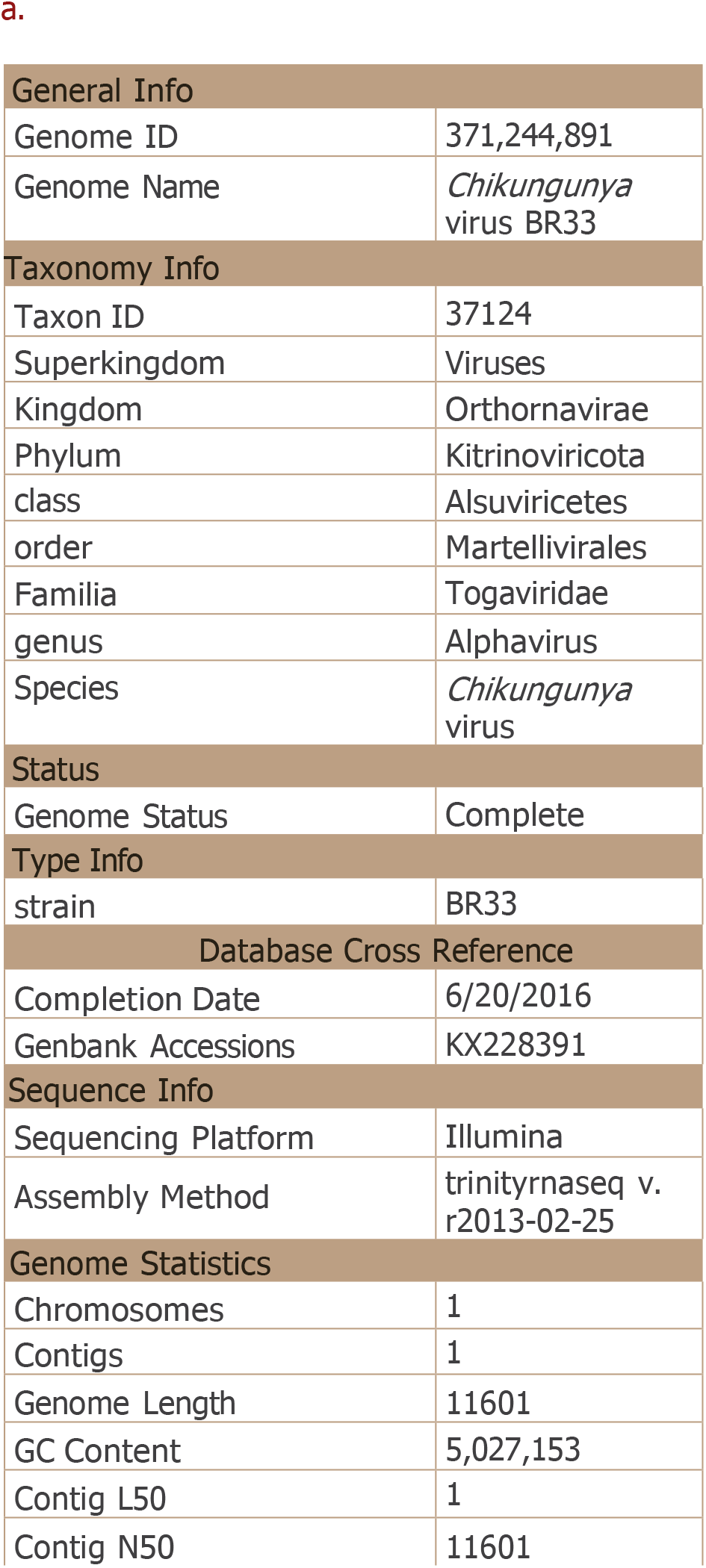
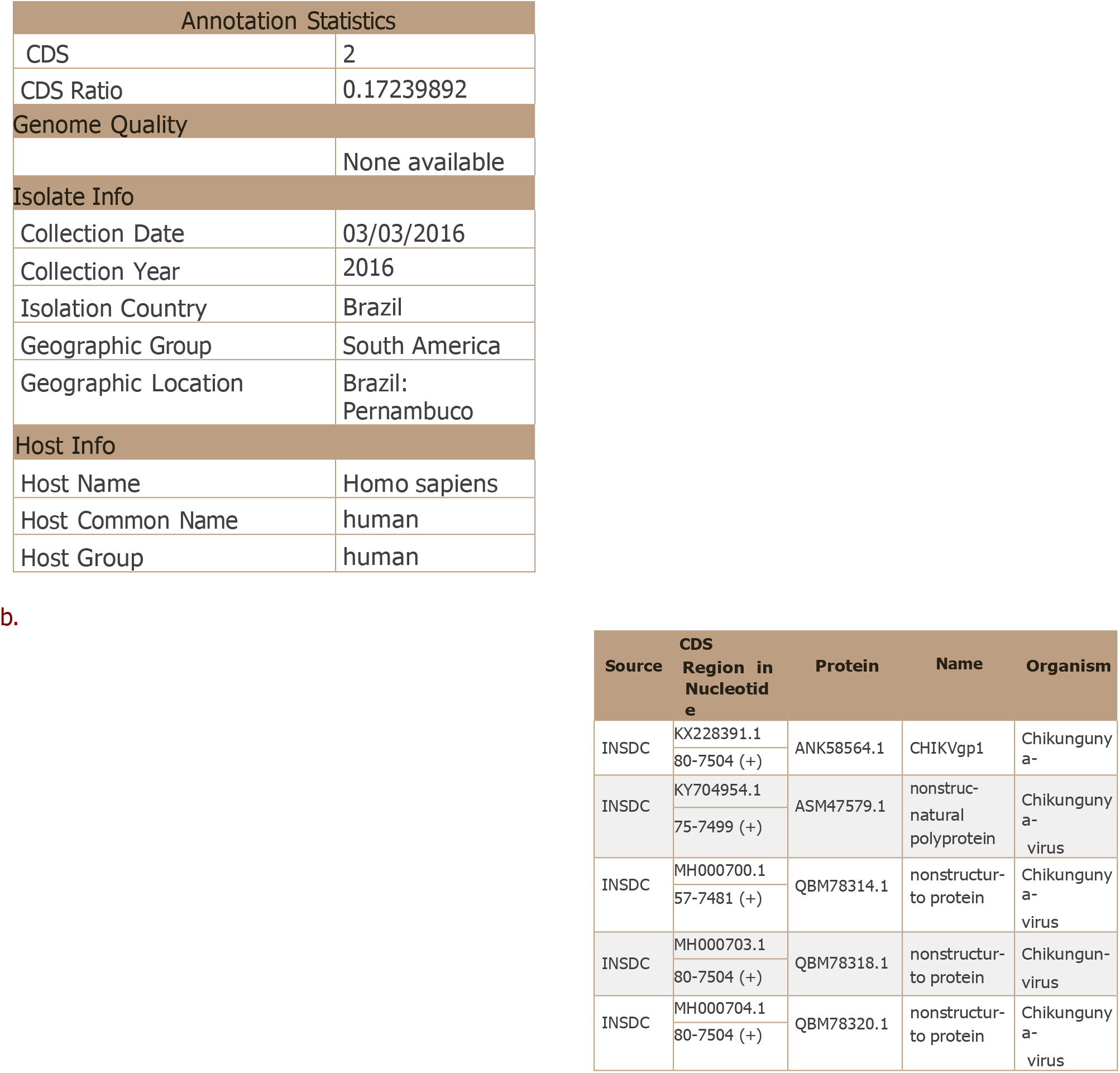
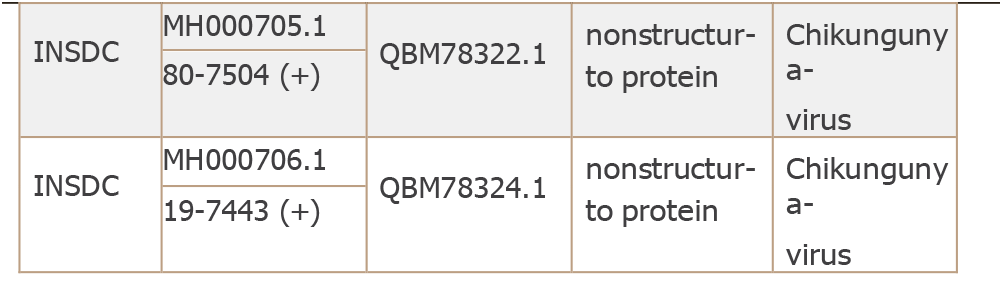
Physical and functional characteristics of CHIKV BR33.

**Figure 1.**
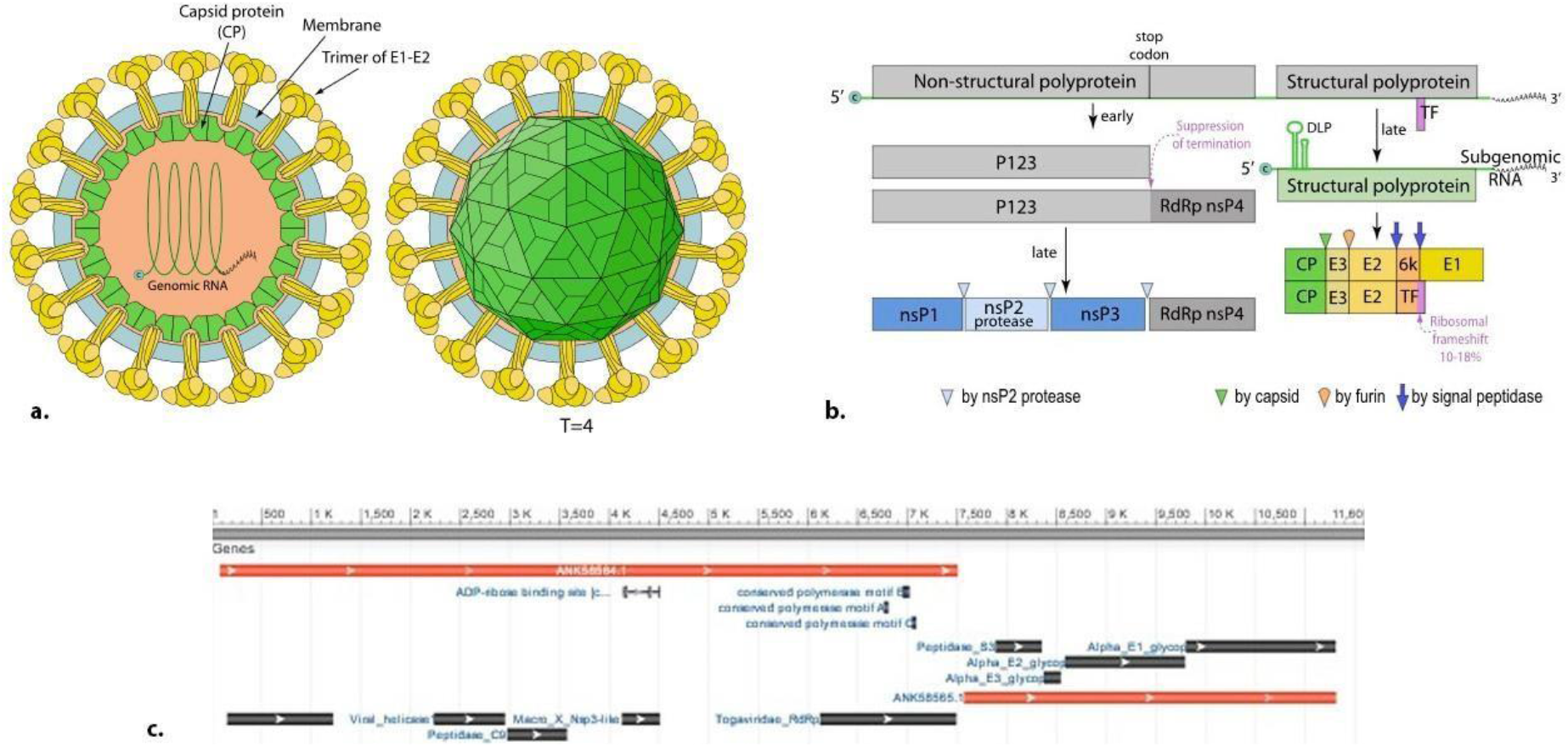
Structural (a) and genetic (b,c) characteristics of CHIKV BR33. Organization of the CHIKV genome and gene products: The genomic organization of *Chikungunya* virus (CHIKV) resembles that of other alphaviruses. It consists of two open reading frames (ORFs), both flanked by structures 5’ cap and a 3’ poly(A) tail. The regions near the 5’ and 3’ ends of the CHIKV genome contain untranslated regions (NTRs), and the junctional region (J) is also noncoding.

**Figure 2.**
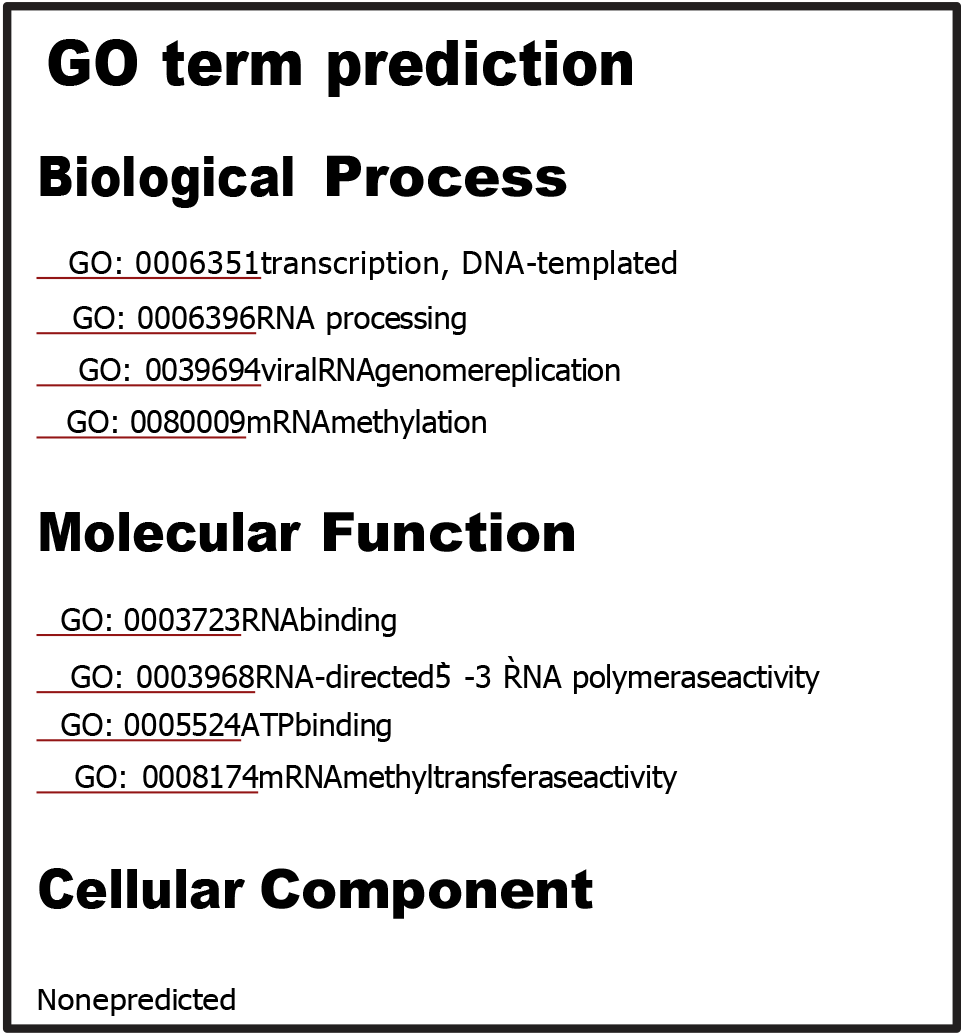
The classification of CHIKgp1 protein by Gene Ontology, divided into functional biological, molecular and cellular components. Gene Ontology (GO): Gene Ontology (GO) constitutes a cornerstone in the field of biological information by offering precise definitions of protein functions. GO is an organized and regulated lexicon consisting of terms known as GO terms. It is classified into three distinct and non-overlapping ontologies: molecular function (MF), biological process (BP) and cellular component (CC). The structure of GO is represented as a directed acyclic graph (DAG), where the terms take the form of nodes and the relationships between terms form the edges. This framework offers greater flexibility than a traditional hierarchy, as each term can have multiple connections to broader parent terms as well as more specific child terms (du Plessis et al. 2011). Genes or proteins are associated with GO terms through an annotation process, which serves as linkage. Each GO annotation has an attributed source and an entry in the databases. These sources can range from bibliographic references to database references and computational evidence. Each biological molecule is linked to the most specific set of terms that accurately describes its functional attributes. Consequently, if a biological molecule is associated with a particular term, it will be inherently connected to all original terms within the hierarchical structure of that term (du Plessis et al. 2011). This comprehensive system enables accurate classification of biological functions and greatly aids in understanding the functionality of genes and proteins.

**Figure 3.**
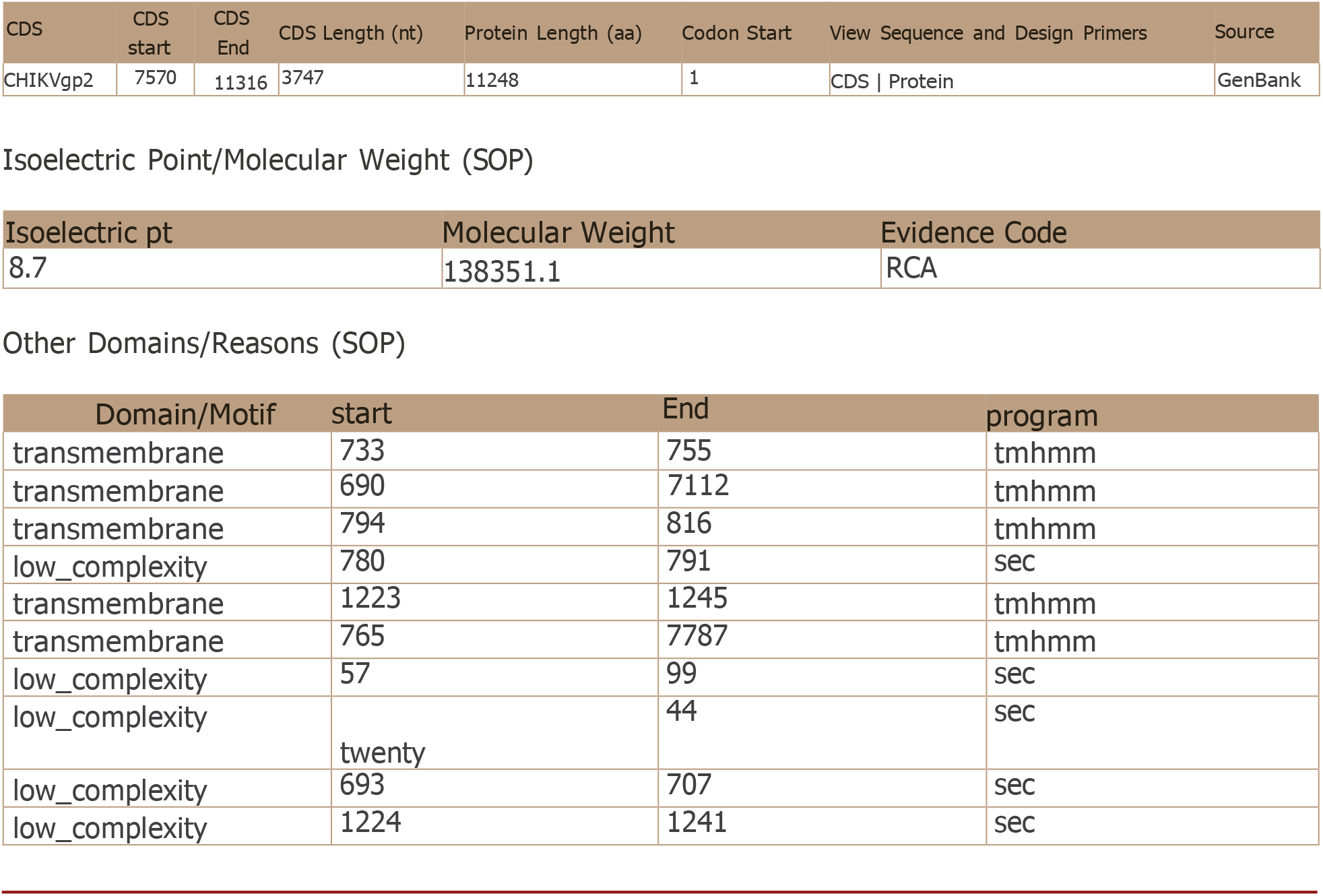
Functional characteristics of annotation and domain search of the CHIKgp2 protein. Sequence annotations provide detailed information about specific regions or features within a protein sequence. These annotations cover a wide range of elements, including post-translational modifications, binding sites, enzyme active sites, local secondary structures, and other features that are reported in the cited references or predicted. Additionally, any discrepancies or conflicts in sequence information between different references are also documented in this way.

**Figure 4.**
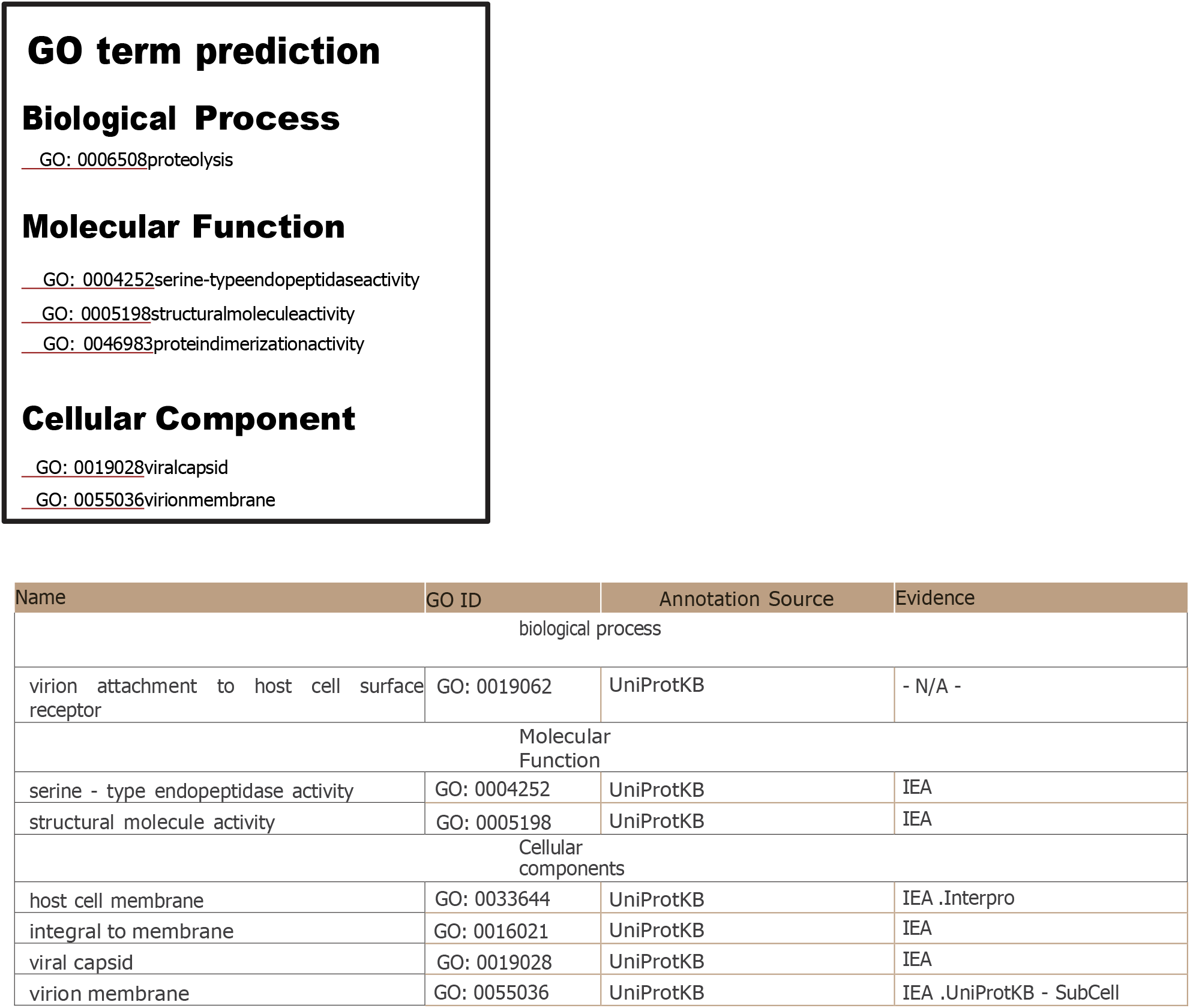
Classification by Gene Ontology of the CHIKgp2 protein, divided into functional biological, molecular and cellular components.

**Figure 5.**
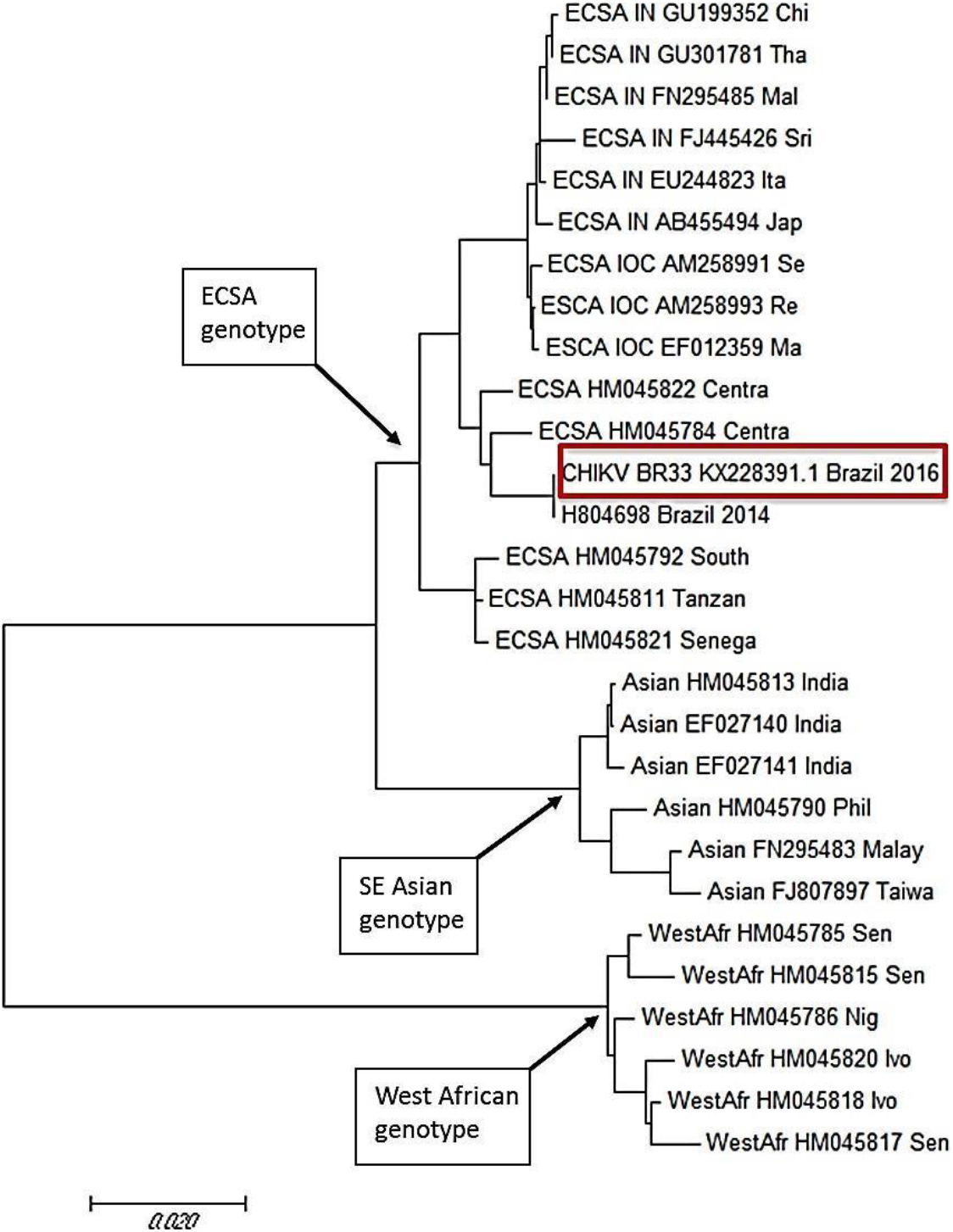
Details of phylogenetic analysis (serotype). CHIKV BR33 Bayesian filament that includes the three genotypes: West Africa (WA), East Central Africa-South Africa (ECSA) and Southeast Asia (SE). The red box shows CHIKV BR33.

The 5’ ORF is directly translated from genomic RNA and encodes four nonstructural proteins (nsP1, nsP2, nsP3, and nsP4). In contrast, the 3’ ORF is translated from a subgenomic 26S RNA and encodes several structural proteins, including the capsid protein (C), two surface envelope glycoproteins (E1 and E2), and two known small peptides. like E3 and 6k. These structural and non-structural proteins, namely nsP1 to nsP4 and C, E1, E2, E3 and 6k, are produced by proteolytic cleavage of polyprotein precursors. This genomic organization and subsequent protein synthesis are fundamental aspects of the CHIKV life cycle and play a crucial role in its pathogenicity and interactions with the host. https://viralzone.expasy.org/625?outline=all_by_species

The results showed that the isolated CHIKV BR33 strain belongs to the ECSA genotype (Fig. 5). A search using the nucleotide search tool, BLASTn (Huh, JE et al. 2021), revealed that the tested strain is closely related to the CHIKV virus strain BHI3734/H804698, with GenBank accession number: KP164568.1 was isolated in Brazil of a patient from Feira de Santana-BA was not presented to the NCBI by the Center for Technological Innovation of the Evandro Chagas Institute. It has a size of 11,812 base pairs and phylogenetically belongs to the ECSA genotype. Other strains isolated by the same research group in 2015, in Feira de Santana, a municipality in the state of Bahía, also located in the Northeast region of Brazil (Nunes et al. 2015), showed a similarity of 99% in terms of nucleotide identity.

## DISCUSSION

CHIKV is an arthropod-borne virus with epidemic potential affecting millions of people worldwide (de Souza WM et al. 2024). The sequencing genomics cobra importance because it allows monitoring the spread of a virus, its changes and how these can affect public health. This is how in this case CHIKV BR33 was diagnosed and identified, showing that the strain belonging to the ECSA genotype is circulating in Brazil. The detection of this CHIKV strain, like the one identified in Rio de Janeiro by Romero Felipe et al. 2023, serves as compelling evidence of its current circulation in Brazil.

The spread of CHIKV between countries and continents is attributed to infected travelers from endemic areas.

This warrants further exploration of its clinical implications, particularly with regard to its virulence and its association with disease severity.

Virulence refers to the ability of the virus to cause disease and the severity of that disease. Understanding the virulence of this strain is crucial as this can significantly affect clinical prognosis. Investigating whether this strain exhibits increased virulence compared to other *Chikungunya* strains is essential to predict the potential public health impact and evaluate the relationship between this strain and disease severity, which is critical. To the best of our knowledge, severity appears to be related to viral genotype, with an overall non-recovery rate of 50% (95% CI 40-60%) in the ECSA/IOL lineage, 36% (95% CI %: 20-52%) associated with the Asian lineage, and 13% with the ECSA lineage (95% CI: 7-18%) (Cunha, M. S et al. 2020, González Velázquez JN et al. 2023) Phase 3 clinical trials in 4,115 adults showing a seroresponse rate of 98.9% at 28 days with a single vaccination, is approved for people over 18 years of age who are at higher risk of exposure to the virus. Serious reactions occurred in 1.6% of recipients; some participants they had reactions adverse prolonged periods that lasted at least 30 days. Even with the existence of a vaccine, knowing genomic sequencing can help in the future determine if virus variants are affecting its effectiveness. This by monitoring the evolution of the virus and detecting the appearance of new variants. Despite the existence of a vaccine, it has not yet been widely distributed, so current treatment depends on symptomatic relief through analgesics, antipyretics, non-steroidal anti-inflammatory drugs and, in cases serious, methotrexate (Bartholomeeusen K et al. 2023).

At the moment No there is medicines Effective antivirals available for the treatment of CHIKV infection. So far the only FDA-approved vaccine for CHIKV is VLA 1553 vaccine, registered as IXCHIQ. Its approval in November 2023 is based on data from The behavior and severity that the symptoms can reach lead to the need to increase efforts to find safe antiviral drugs against CHIKV. The use of antivirals during acute infection could reduce the likelihood of developing chronic symptoms. The proteins nsP2 (protease), nsP4 (viral polymerase), and Mxra8 have been studied as targets of antiviral drugs that work for arthritogenic alphaviruses such as CHIKV. Sequencing is expected to help evaluate the effectiveness of these treatments in each genotype (Abdelnabi R et al. 2020).

Furthermore, the problem is not only limited to Brazilian territory because there is a new circulating strain that can increase the virulence of CHIKV, it can also increase the reach and spread of the virus throughout the entire Latin American territory, which is why it was completely necessary to know genetically to this new strain to find alternatives for mitigation and control. As we can see in the reports from the ministries of health, although dengue continues to be the main arbovirus in America, cases of *Chikungunya* are also increasing, warranting both epidemiological and clinical control.

Measures to ensure adequate clinical management of suspected arbovirus cases should be a priority. Capacities must be strengthened at the primary health care level and from this level prevent progression to severe forms and deaths. This requires early clinical diagnosis and recognition of warning signs (such as intense and sustained abdominal pain or pain on palpation of the abdomen, persistent vomiting, clinical fluid accumulation, mucosal bleeding, lethargy, restlessness, enlargement of the liver > 2 cm below the costal margin and progressive increase in hematocrit) in order to initiate appropriate management according to the recommendations published in the clinical guidelines.

Laboratory results should be analyzed with clinical information and according to epidemiological context, for surveillance and not for clinical decision making.

Laboratory confirmation of infection is based on virological (RT-PCR and ELISA, and in some cases viral isolation in culture for additional characterization) and serological (IgM detection) tests. However, to confirm cases, virological tests that demonstrate the presence of the complete virus, its genetic material or its proteins should be prioritized. In fatal cases, tissue samples (liver, spleen, kidney) should be considered both for the detection of genetic material (RT-PCR) and for histopathological and immunohistochemical study. Taking biopsies in a patient with suspected *Chikungunya* is completely contraindicated. Since laboratory services are a key component of CHIKV epidemiological and virological surveillance, timely detection and characterization in appropriate samples must be maintained. As far as possible and according to the capabilities of each laboratory, it is recommended to take samples from 100% of serious cases and fatal, while only a proportion (10-30% or a maximum number of samples depending on installed capacity) of those cases without alarm signs will be necessary for surveillance. The present study helps to better understand the pathophysiology and molecular aspects of the disease by achieving a genetic and functional description complete of the virus and its structural proteins.

In summary, the detection of this specific strain of *Chikungunya* in Brazil warrants a deeper exploration of its clinical implications, particularly in relation to its virulence and its association with the severity of the disease. Furthermore, it is essential to evaluate the relationship between this strain and the severity of the disease. Some strains of *Chikungunya* have been associated with more severe clinical manifestations, such as a higher incidence of severe joint pain, arthritis, and complications. neurological. If this African strain shows a stronger correlation with severe disease could have important implications for health systems and clinical management strategies in regions affected by the virus.

Furthermore, the genetic diversity of the virus could potentially influence the effectiveness of diagnostic tests, treatment approaches and vaccine development. An exhaustive study of the specific genetic characteristics of this strain is warranted to evaluate these aspects in depth. By delving into the clinical implications, we can gain insight valuable information on the potential impact of this African strain of *Chikungunya* on the health of affected populations and, consequently, inform more specific and effective public health interventions and clinical management strategies (Lima-Camara, 2016).; Zerfu et al. 2023).

Thus, evidence such as that presented in this document is crucial for the alert and continuous surveillance of national and international disease control organizations, for the prevention of new cases that could collapse health services during the simultaneous explosive epidemics that circulate in all Latin American countries (Lima-Cámara, 2016; Zerfu et al. 2023). It is important to phylogenetically identify the genotype circulating in the Brazilian outbreak, because the presence of genotypes such as ECSA has also been associated with an increase in mosquito infectivity, followed by Asian strains (Hakim & Aman, 2022; Kumar et al. 2014; For these reasons, the present study helps to better understand the pathophysiology and molecular aspects of the disease, achieving a complete genetic and functional description of the virus and its structural proteins, relating this to the virulence capacity presented by this viral species with epidemic potential. In this way the complete *Chikungunya* genome generated in this research (GenBank: KX228391.1) It has been fundamental in multiple investigations worldwide, for the creation of diagnostic tests, descriptive phylogenetic trees, creation of vaccines among other research developments (Tuekprakhon A et al. 2018, Machado LC et al. 2019, Laurence Thirion et al. 2019, Reddy A et al. 2020, Su Qianling et al.2019).

## CONCLUSION

To our knowledge, this is the first report of a complete genome of CHIKV, isolated in Brazil, in 2016. The described sequence, additional phylogenetic analyzes of these genomes and the other sequences will be detailed in future publications. Sequencing the complete of the spread of the virus in different regions, which is essential for the implementation of control and prevention measures, especially in areas where the virus is endemic. Knowledge of the genetics of the *Chikungunya* virus is crucial for development of treatments genome of the *Chikungunya* virus is of utmost antivirals effective. These importance for several reasons. First of all, it allows different strains and mutations of the virus to be identified. This is crucial to Drugs can target specific components of the viral genome to inhibit its replication.

Understanding the genetic diversity of the virus and its ability to evolve, allowing sequencing Finally, genomics the us effective surveillance of mutations, which is vital to predict the spread of the virus and develop more effective prevention and treatment strategies. In addition, genomic sequencing facilitates diagnosis and the development of specific tests. These tests play a fundamental role in detecting the presence of the virus in infected patients, allowing quick and accurate decisions to be made about control measures. Furthermore, this genomic information is essential for designing vaccines against *Chikungunya*. A detailed understanding of the viral genome allows the creation of vaccines capable of eliciting effective and specific immune responses against the virus. The sequencing genomics also provides valuable information for epidemiological studies and understanding allows monitoring the possible emergence of drug resistance antivirals used in the treatment of *Chikungunya*, which is essential to adjust the treatments and ensure their effectiveness over time. In summary, sequencing the *Chikungunya* virus genome is a critical component in fighting this disease and safeguarding public health.

## Acknowledgment

The authors are especially grateful to Dr. Irene Bosch of the Columbia Mailman School of Public Health, New York, NY, USA and Dr. Lee Gehrke, director of the Gehrke Laboratory at HARVARD MEDICAL SCHOOL (HMS), Boston, Massachusetts, USA and the Massachusetts Institute of Technology (MIT), Cambridge, MA, USA for the academic training, methodological advice and loan of equipment to carry out this research.

## AUTHOR CONTRIBUTIONS

RGVJ designed the study; collected and prepared the biological materials; performed the experiments; performed data analysis; drafted and wrote the manuscript. The other authors contributed to the writing, edited, read and approved the final draft.

